# Simultaneous single-cell profiling of lineages and cell types in the vertebrate brain by scGESTALT

**DOI:** 10.1101/205534

**Authors:** Bushra Raj, Daniel E. Wagner, Aaron McKenna, Shristi Pandey, Allon M. Klein, Jay Shendure, James A. Gagnon, Alexander F. Schier

## Abstract

Hundreds of cell types are generated during development, but their lineage relationships are largely elusive. Here we report a technology, scGESTALT, which combines cell type identification by single-cell RNA sequencing with lineage recording by cumulative barcode editing. We sequenced ~60,000 transcriptomes from the juvenile zebrafish brain and identified more than 100 cell types and marker genes. We engineered an inducible system that combines early and late barcode editing and isolated thousands of single-cell transcriptomes and their associated barcodes. The large diversity of edited barcodes and cell types enabled the generation of lineage trees with hundreds of branches. Inspection of lineage trajectories identified restrictions at the level of cell types and brain regions and helped uncover gene expression cascades during differentiation. These results establish scGESTALT as a new and widely applicable tool to simultaneously characterize the molecular identities and lineage histories of thousands of cells during development and disease.

## INTRODUCTION

Recent advances in single-cell genomics have spurred the characterization of molecular states and cell identities at unprecedented resolution^1–3^. Droplet microfluidics, multiplexed nanowell arrays and combinatorial indexing all provide powerful approaches to profile the molecular landscapes of tens of thousands of individual cells in a time-and cost-efficient manner^4–8^. Single-cell RNA sequencing (scRNA-seq) can be used to classify cells into “types” using gene expression signatures, and generate catalogs of cell identities across tissues. Such studies have identified novel marker genes and revealed cell types that were missed in prior bulk analyses^9–15^

Despite this progress, it has been challenging to determine the developmental trajectories and lineage relationships of cells defined by scRNA-seq. The reconstruction of developmental trajectories from scRNA-seq data requires deep sampling of intermediate cell types and states^16–20^ and is unable to capture the lineage relationships of cells. Conversely, lineage tracing methods using viral DNA barcodes, multicolor fluorescent reporters or somatic mutations have not been coupled to single-cell transcriptome readouts, hampering the simultaneous large-scale characterization of cell types and lineage relationships^21,22^.

Theoretically, such limitations could be overcome if lineage and cell type information were both encoded in a cell’s transcriptome. Here we implement this strategy by combining scRNA-seq with GESTALT^23^, one of several lineage recording technologies based on CRISPR-Cas9 editing^24–28^. In GESTALT, the combinatorial and cumulative addition of Cas9-induced mutations in a genomic barcode creates diverse genetic records of cellular lineage relationships. Mutated barcodes are sequenced, and cell lineages are reconstructed using tools adapted from phylogenetics^23^. We demonstrated the power of GESTALT for large-scale lineage tracing and clonal analysis in zebrafish but encountered two limitations. First, edited barcodes were sequenced from genomic DNA of dissected organs, resulting in the loss of cell type information. Second, barcode editing was restricted to early embryogenesis, hindering reconstruction of later lineage relationships. Here we overcome these limitations: we used scRNA-seq to simultaneously recover the cellular transcriptome and the edited barcode expressed from a transgene, and we created an inducible system to introduce barcode edits at later stages of development (Fig. 1). Application of this technology, scGESTALT, to the zebrafish brain identified more than 100 different cell types and created lineage trees that helped reveal spatial restrictions, lineage relationships, and differentiation trajectories during brain development. The building blocks and methods underlying scGESTALT are transferable to many multicellular systems, making it a widely applicable technology to simultaneously uncover cell type and lineage for thousands of cells.

**Figure 1.**
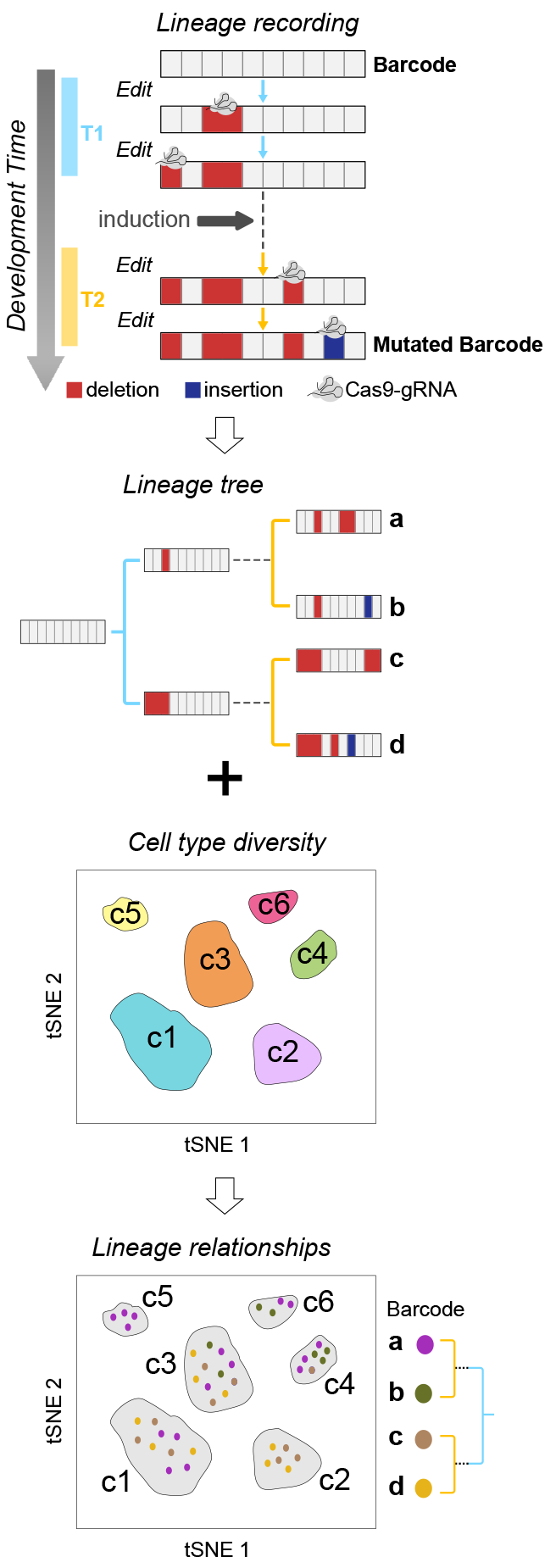
scGESTALT: Simultaneous recovery of transcriptomes and lineage recordings from single cells. During development, CRISPR-Cas9 edits record cell lineage in mutated barcodes (a,b,c,d). Barcode editing occurs at early (T1, blue) and late (T2, yellow) timepoints during development. Simultaneous recovery of transcriptomes and barcodes from the same cells can be used to generate cell lineage trees and also classify them into discrete cell types (c1 - c6).

## RESULTS

### Droplet scRNA-seq identifies cell types and marker genes in the zebrafish brain

To identify cell types in the zebrafish brain with single-cell resolution, we dissected and dissociated brains from 23-25 days post-fertilization (dpf) animals (corresponding to juvenile stage) and encapsulated cells using inDrops^4^ (Fig. 2a and Supplementary Fig. 1). We used manually dissected whole brains and forebrain, midbrain and hindbrain regions. In total, we sequenced the transcriptomes of ~66,000 cells with an average of ~22,500 mapped reads per cell. After filtering out lower quality libraries, we generated a digital gene expression matrix comprising 58,492 cells with an average of ~3,100 detected unique transcripts from ~1,300 detected genes per cell. We used an unsupervised, modularity-based clustering approach^5,29^ to group all cells into clusters (Fig. 2b) and initially identified 63 transcriptionally distinct populations. All clusters were supported by cells from multiple biological replicates.

To classify each cluster, we systematically compared differentially expressed genes with prior annotations of gene expression in specific cell types or brain regions in the literature and the ZFIN database^30,31^. Initial analysis identified 45 neuronal subtypes, 9 neural progenitor classes, 3 oligodendrocyte clusters, microglial cells, ependymal cells, blood cells and vascular endothelial cells (Supplementary Fig. 2 and Supplementary Table 1). We were able to resolve all but three neuronal clusters (clusters 0, 24 and 31), with cluster 0 likely corresponding to nascent neurons mostly from the forebrain, as it displays high levels of *tubb5* expression and moderate levels of *neurod 1* and *eomesa*. We captured multiple cell types that each comprised less than 1% of all profiled cells. These include *aanat2*^+^ neurons from the pineal gland (cluster 62), representing 0.04% of captured cells; *sst1.1*^+^ and *npy*^+^ neurons in the ventral forebrain (cluster 53, 0.34% of data); *aldoca*^+^ Purkinje neurons in the cerebellum (cluster 43, 0.65% of data); and fluorescent granular perithelial cells (cluster 54, 0. 33% of data), a population of perivascular cells recently described in zebrafish^32^. Using marker genes and gross spatial information from manually dissected brain regions, most clusters could be assigned to specific brain regions (e.g. hypothalamus in forebrain and cerebellum in hindbrain) (Fig. 2c and Supplementary Fig. 1). Spatially restricted transcription factors were enriched in specific clusters, including *dlx2a, dlx5a, emx3* and *foxg1a* in forebrain clusters; *barhl2, gata2a, otx2*, and *tfap2e* in midbrain clusters; and *phox2a, phox2bb*, and *hoxb3a* in hindbrain clusters. Thus, regional location in the brain was a strong contributor to gene expression differences and drove clustering outcomes.

To identify diverse cell types that might have been masked when analyzing the whole dataset in bulk, we performed a second round of clustering on the larger neuronal clusters (Fig. 2d, Supplementary Fig. 3 and Supplementary Table 2). For example, reanalysis of the eight initial hindbrain and cerebellum clusters identified 17 transcriptionally distinct groups (Fig. 2e). After removing five subclusters that did not separate further from the original clusters or had no clear gene markers, we classified the 12 remaining subclusters. For example, cluster 23 (hindbrain) split into three subclusters enriched in *hoxb3a* (s9), *hoxb5b* (s10) and *pou4f1* (s15). Combined with the whole-dataset clustering results, iterative analyses identified a total of 102 transcriptionally distinct cell types in the brain.

A large subset of sequenced cells (~13%, 8 clusters) was composed of neural progenitors (Fig. 2b), consistent with the continuous growth and neurogenesis in the zebrafish brain^33^. The distinct categories of progenitor clusters corresponded to quiescent radial glia cells, which are the neural stem cells of the brain and express *gfap, fabp7a* and *s100b* (clusters 25, 33, 48); intermediate progenitors expressing proneural transcription factors such as *ascl1a, neurog1* and *insm1a* (8, 17); and highly proliferative progenitors expressing *pcna, mki67* and *top2a* (clusters 19, 22, 44) (Fig. 2f). Although three progenitor clusters could be assigned to specific regions, gene expression profiles suggested that most progenitors were more closely related to other progenitors than to their differentiated neighbors (Fig. 2c).

**Figure 2.**
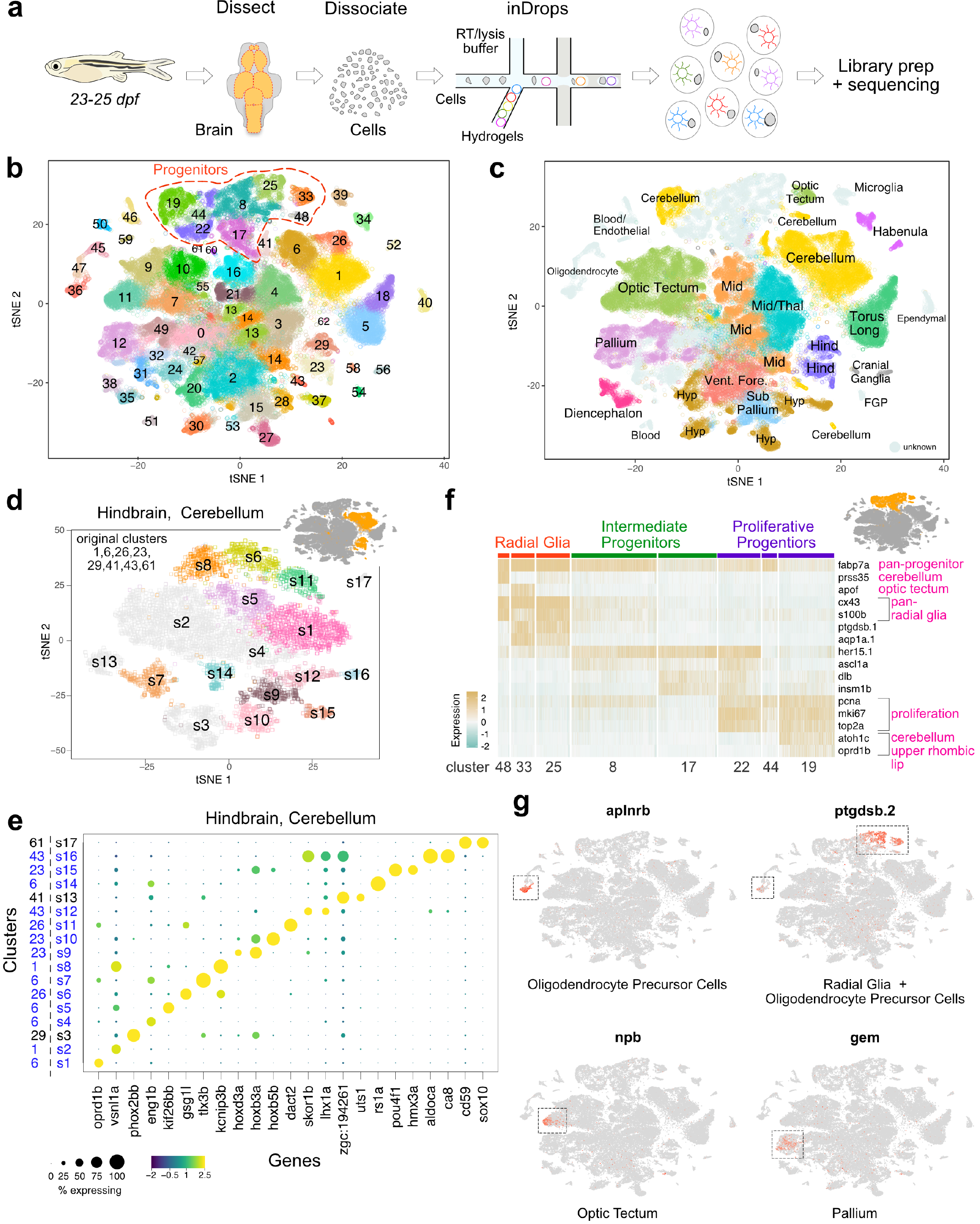
Cell type diversity in the juvenile zebrafish brain. **a**. Juvenile zebrafish brains were dissected, dissociated and processed by inDrops. **b**. t-SNE plot of 58,492 cells clustered into 63 cell types. Progenitor cell types highlighted. **c**. t-SNE plot with cell clusters labeled with inferred anatomical location. FGP, fluorescent granular perithelial cells. Hind, hindbrain. Hyp, hypothalamus/preoptic area. Mid, midbrain. Thal, thalamus. Torus Long, torus longitudinalis. Vent. Fore., ventral forebrain. Cells of unknown origin or broad distribution are colored in grey. **d**. Iterative clustering of cells from the hindbrain/cerebellum. Inset highlights these eight clusters within initial t-SNE plot. Main panel, t-SNE plot of the resulting subclusters. Subclusters colored light grey either did not partition further or had no clear markers. **e**. Dotplot of gene expression patterns of select marker genes (columns) for each subcluster (rows) from the hindbrain/cerebellum. Dot size represents the percentage of cells expressing the marker; color represents the average scaled expression level. Initial cluster numbers are indicated to the left of the subcluster (s) number. Clusters colored blue were subdivided by iterative analysis. **f**. Heat map of scaled gene expression of representative marker genes across cells within eight neural progenitor clusters. Original cluster numbers are indicated on the bottom. Marker genes are categorized according to the cell types they label (pink text). Inset highlights these eight clusters within initial t-SNE plot. **g**. Gene expression patterns of novel cell type markers. Cells within each t-SNE plot are colored by marker gene expression level (grey is low, red is high). Dotted boxes highlight clusters where markers are enriched.

Differential gene expression identified previously unrecognized marker genes (Fig. 2g). For example, *aplnra* and *aplnrb*, G-protein-coupled receptors that are involved in cell migration^34^, were highly enriched in oligodendrocyte precursor cells (OPC). Subpopulations of quiescent and dividing radial glia cells, as well as OPCs, expressed *ptgdsb.1* and *ptgdsb.2*, enzymes that regulate synthesis of prostaglandin D2. *Npb* (neuropeptide b) and *gem* (GTP binding protein overexpressed in skeletal muscle) transcripts were detected in subclusters of optic tectum and pallium cells, respectively.

Taken together, these results provide the first global catalogue of progenitor and mature cell types in the zebrafish brain and provide a resource for the study of specific cell populations and marker genes in a vertebrate brain.

### Inducible Cas9 expression enables late barcode editing

Neurogenesis occurs after the onset of gastrulation, making lineage trajectories in the brain most informative after this developmental stage. In our initial implementation of GESTALT, all editing reagents (Cas9 protein and sgRNAs) were injected into one-cell stage embryos, thus centering barcode editing on pre-gastrulation stages^23^. To overcome this limitation and enable recording of lineages at later stages, we generated transgenic zebrafish wherein Cas9 activity could be induced in vivo using a promoter activated by heat shock and a subset of sgRNAs (sgRNAs 5-9) were constitutively expressed via U6 promoters. We then devised a new strategy whereby editing activity was split across two time windows (Fig. 3a): we crossed the GESTALT barcode transgenic to the inducible Cas9 transgenic and injected single-cell embryos with Cas9 protein and sgRNAs 1-4. This strategy initiates an “early” round of Cas9 activity that edits barcodes in embryonic cells at target sites 1-4. We then heat shocked the embryos at 30 hours post-fertilization (hpf) to induce ubiquitous expression of transgenic Cas9. This strategy resulted in a “late” round of Cas9 activity that used the zygotically expressed sgRNAs to edit barcodes at sites 5-9. To evaluate this two-timepoint editing strategy, we extracted genomic DNA from 55 hpf control and edited double transgenic embryos, and amplified and sequenced GESTALT barcodes^23^. We observed no substantial editing of the barcode when Cas9 and sgRNAs were not injected or expressed in the embryo (Fig. 3b). Upon heat shock, mutations were predominantly confined to sites 5-9 of the barcode. As expected, after injection and subsequent heat shock, barcodes contained edits in “early” sites 1-4 and “late” sites 5-9. We found that all recovered barcodes were edited (100% editing frequency) with a median of 4 independent edits per barcode. Each embryo had a median of 1,504 distinct barcodes (range 731 to 2,213), demonstrating the efficiency of the editing strategy for generating barcode diversity.

**Figure 3.**
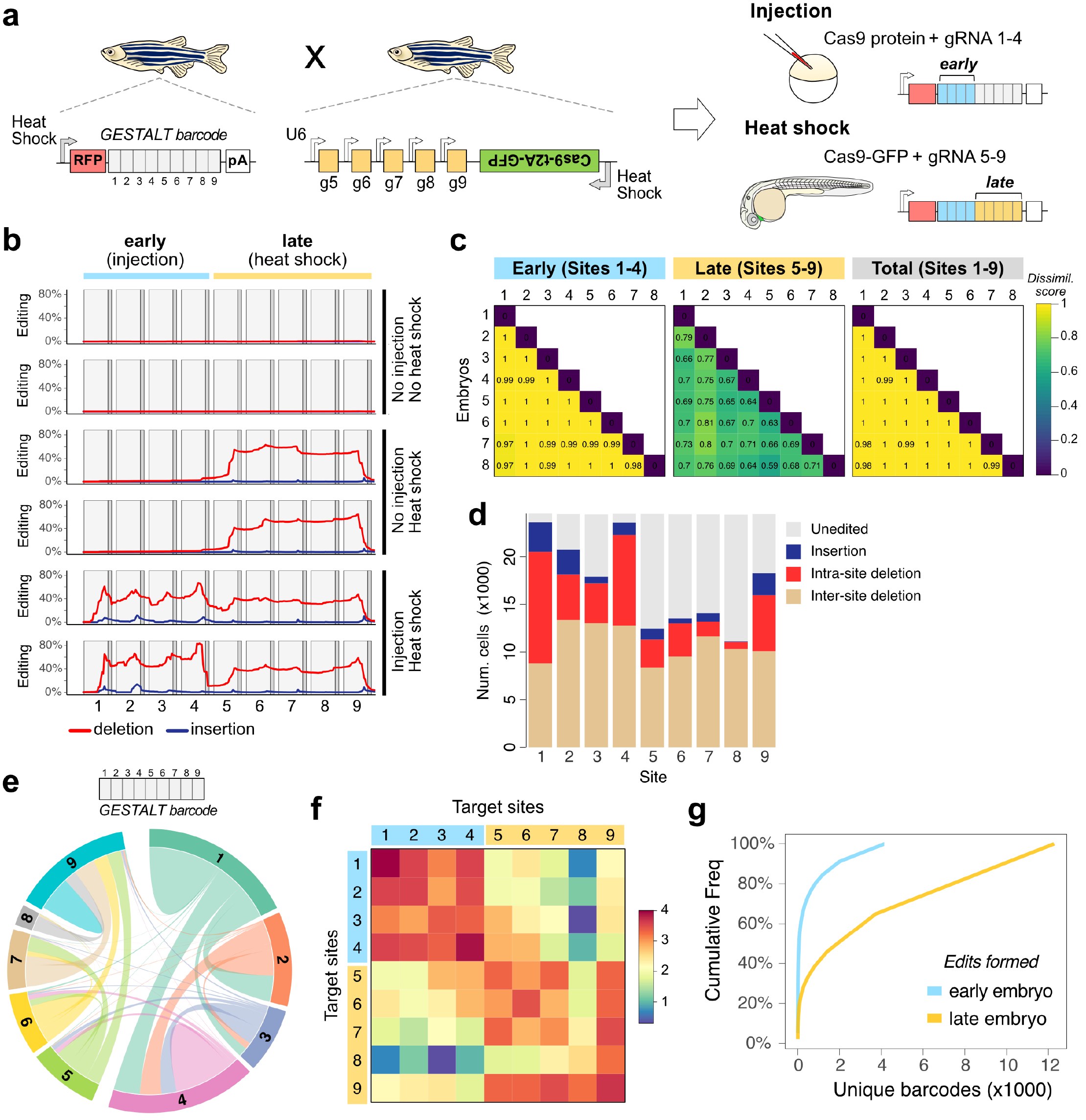
An inducible CRISPR-Cas9 system for late barcode editing. **a**. Zebrafish that express the GESTALT barcode as polyadenylated (pA) mRNA were crossed to zebrafish that express heat shock-inducible Cas9 along with gRNAs 5-9. Resulting embryos were injected with Cas9 and gRNAs 1-4 at the one-cell stage (blue bars; early editing), and heat shocked at 30 hpf to induce transgenic Cas9 for a second round of editing (yellow bars; late editing). **b**. Mutations within the nine CRISPR target sites of the GESTALT barcode for three editing conditions (2 animals per condition). Red lines represent deletions, blue lines represent insertions. **c**. Pairwise comparisons using cosine dissimilarity of early and late edit patterns from eight doubly-edited embryos. **d**. Edit type at each target site within the barcode from all eight doubly-edited embryos. **e**. Chord diagram of the nature and frequency of deletions within and between target sites. Each colored sector represents a target site. Links between target sites represent intersite deletions; self-links represent intra-site deletions. Link widths are proportional to the edit frequencies. **f**. Heat map of the frequency (log10 scale) of inter-site and intra-site deletions within and across the barcode target sites. **g**. Cumulative frequency of each barcode across all cells pooled from 8 embryos, considering only early barcode edits (blue) and full barcodes (yellow).

To quantify the diversity of barcodes resulting from two-timepoint editing, we compared editing outcomes in different embryos (n = 8). Only 63 of the 12,277 distinctly edited barcodes (0.5%) were present in more than one embryo, demonstrating highly independent generation of barcodes (Fig. 3c). To assess the spectrum of barcode repair products, we profiled the nature (insertion, deletion) and frequency of edits within all 24,360 recovered barcodes. The landscape of intra-site (edits within a site) and inter-site (edits that span two or more sites) deletions varied highly among the different target sites, revealing a large “sequence space” available for DNA repair outcomes from two-timepoint editing (Fig. 3d-f and Supplementary Fig. 4).

The addition of late edits to earlier edits during two-timepoint editing predicts increased barcode diversity. Indeed, full barcodes containing both early and late edits were higher in number and less clonal compared to the early edited barcodes (Fig 3g). 4,141 early barcodes diversified to 12,277 full barcodes. Each early barcode was observed in on average 2.97 distinct late barcodes (range 1 to 534). The diversity and editing efficiency was higher in the early sites as compared to the late sites (Fig. 3b, c). Later edits also resulted in more inter-site deletions. This difference might reflect the activity of distinct DNA repair pathways^35,36^ during development or susceptibility to re-cleavage from the extended presence of Cas9-sgRNA ribonucleoprotein during slower cell cycles at later stages. Collectively, these results show that Cas9-mediated editing is inducible at later stages of development, and in combination with early editing generates thousands of different barcodes.

### scRNA-seq simultaneously recovers single-cell transcriptomes and lineage barcodes

To implement our goal of embedding both lineage and cell type information in a cell’s transcriptome, we introduced the barcode into the 3’ UTR of a heat shock-inducible DsRed transgene (Fig. 3a). Upon heat shock, the edited barcode is expressed as part of the DsRed mRNA and can be isolated with the cellular transcriptome. To test this technology (scGESTALT), we performed two-timepoint editing at the one-cell stage and at 30 hpf and dissected whole brains at 23-25 dpf. Single cells were processed by inDrops, enabling hybridization of endogenous mRNAs and lineage barcode mRNAs to oligodT primers on hydrogels. Barcode libraries were prepared by PCR enrichment of lineage barcode cDNAs (see Methods) and sequenced, resulting in barcode recovery from 3,731 cells from three (ZF1, ZF2, ZF3) juvenile zebrafish brains (750, 2,605 and 376 cells; corresponding to 6%-28% of all profiled cells per animal). To test if barcode recovery might be biased to specific cell types, we compared the cell types identified by scRNA-seq with the identity of cells with recovered barcodes. Strikingly, scGESTALT barcodes overlapped nearly all broadly defined cell types (62/63 broad clusters), indicating that the lineage transgene is widely expressed in the brain. We obtained a range of 150 to 342 distinct barcodes per animal, with a median of 1 (ZF1 and ZF3) or 3 (ZF2) cells, and found no shared barcodes between animals. These results establish scGESTALT as a technology that enables the simultaneous recovery of edited barcodes and transcriptomes from single cells.

### Reconstructed lineage trees reveal relationships between neural cell types

To determine if scGESTALT can reveal lineage relationships, we reconstructed lineage trees for the recovered barcodes using a maximum parsimony approach (see Methods) that anchored the tree with edits at sites 1-4 and extended it with edits at sites 5-9. scGESTALT generated highly branched multi-clade lineage trees. For example, the ZF1 and ZF3 lineage trees comprised 25 and 23 major clades (marked by at least one early edit) that diversified into 193 and 150 late nodes with 341 and 256 branches, respectively (Fig. 4 and Supplementary Fig. 5). Most late edits defined a single node branching from an earlier-marked node, but we also detected as many as 24 late nodes branching from an early-marked node. Thus, late edits greatly increased the branching of the lineage tree. These results provide the proof-of-concept that scGESTALT can reconstruct lineage trees from single-cell transcriptomes.

**Figure 4.**
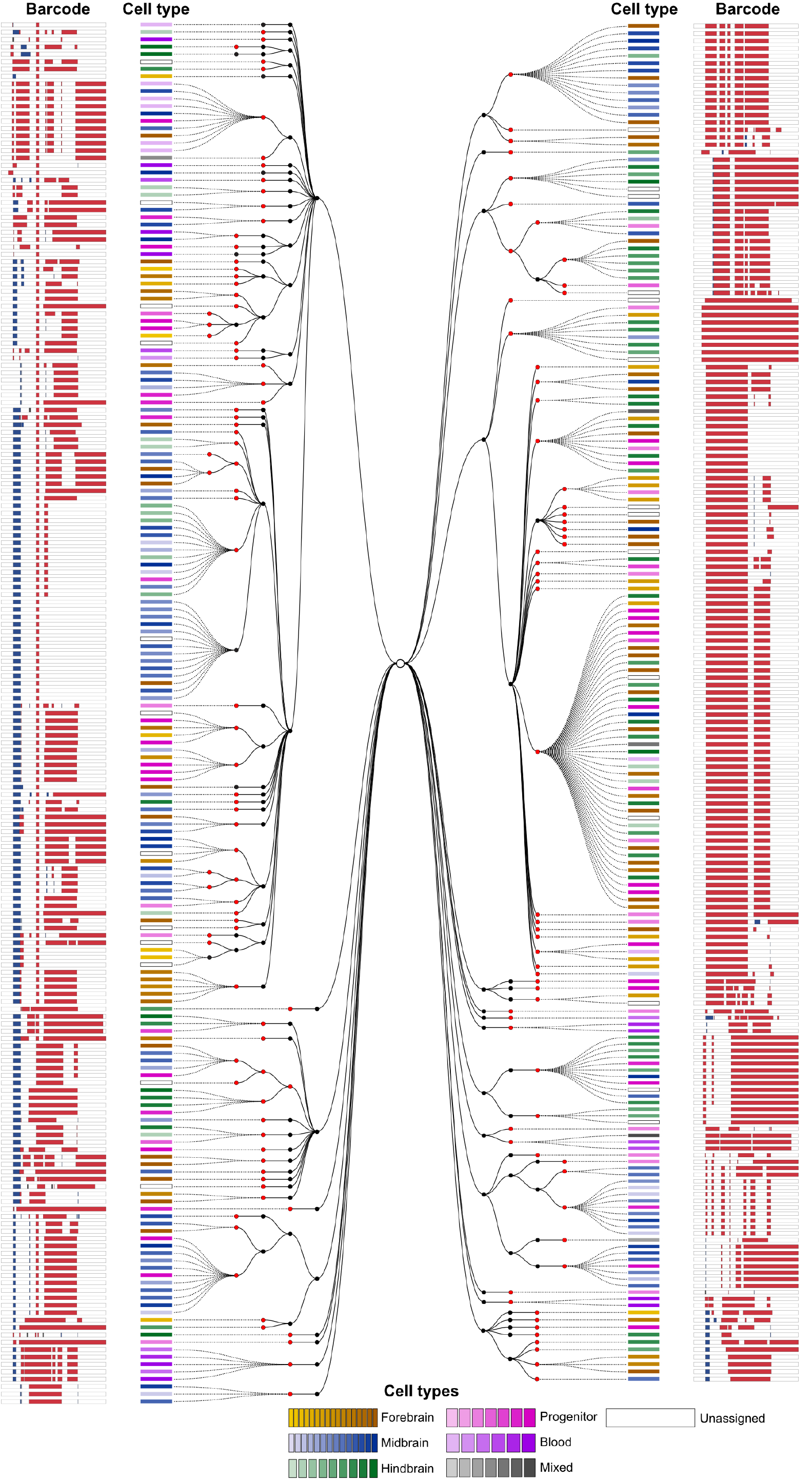
A lineage tree of a zebrafish brain generated using scGESTALT. 376 barcodes recovered from a single juvenile zebrafish brain (ZF3) using scRNAseq were assembled into a cell lineage tree based on shared edits using a maximum parsimony approach. Black nodes indicate early barcode edits; red nodes indicate late edits. Dashed lines connect individual cells to nodes on the tree. Cell types (identified from simultaneous transcriptome capture) are color coded as indicated in the legend. The barcode for each cell is displayed as a white bar with deletions (red) and insertions (blue). A lineage tree for ZF1 is shown as Supplementary Figure 5. Interactive trees and the very large lineage tree for ZF2 can be found at: http://krishna.gs.washington.edu/content/members/aaron/fate_map/harvard_temp_trees/

**Figure 5.**
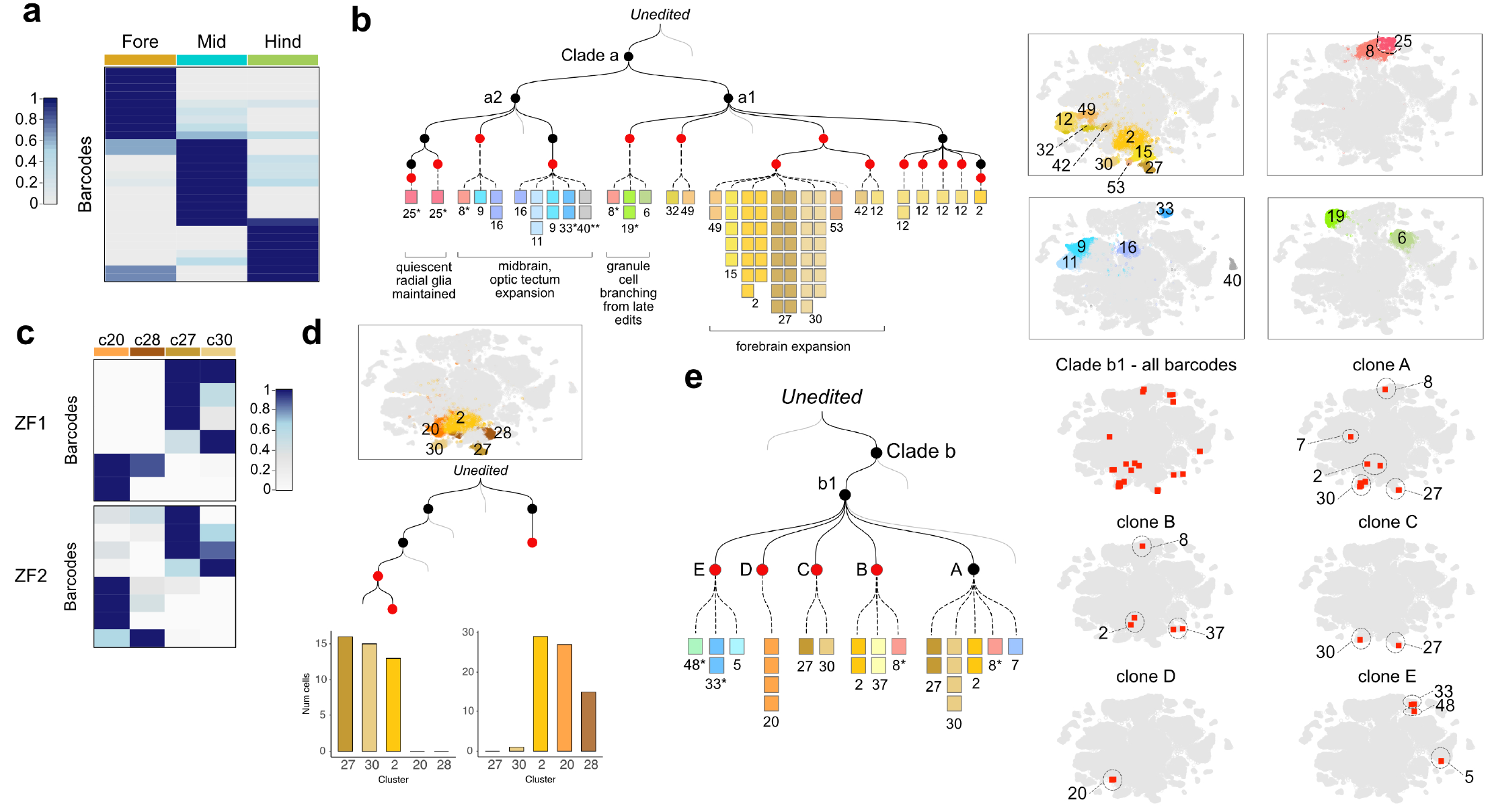
Lineage relationships of cell types in the juvenile zebrafish brain. **a**. Heat map of the distribution of ZF1 barcodes (rows, clone size >= 4 cells) for each region of the brain (columns). Cell types were classified as belonging to the forebrain, midbrain or hindbrain, and the proportions of cells within each region were calculated for each barcode. Region proportions were scaled by row and colored as shown in the legend. **b**. Mini tree showing lineage branches and cluster contributions from clade a within brain ZF1. Black nodes indicate early edits; Red nodes, late edits. Each square represents a cell colored by cell type. Right, t-SNE plots with highlighted cell types: Yellow/brown (forebrain), blue (midbrain), green (hindbrain). Asterisk, progenitor cell types. Double asterisk, ependymal cells. Grey lines, additional branches of the tree. **c**. Heat map of the distribution of ZF1 (6 barcodes, 95 cells) and ZF2 barcodes (8 barcodes, 113 cells) across indicated cell types within the hypothalamus/preoptic area, plotted as above. Insufficient recovery of barcodes from these cell types in ZF3 precluded analysis. **d**. Bar plots showing the distribution of descendant cells from two ZF1 barcodes into cell types of the hypothalamus/preoptic area. **e**. Mini tree showing ZF1 clade b descendants. Subclade b1 was marked during the early round of editing. Clones A, B, C and D were marked during the late round. Clone E was not edited in the late round. The mini tree highlights branches where cluster 20 cells (D) separated from clusters 27 and 30 cells (C) during late barcode editing. Right, t-SNE plots showing barcode distributions across cell types.

To determine the relationship of cells with respect to their cell type and position, we inspected the tree vis-à-vis the identity of cells. Analysis of groups of 4 or more cells with the same barcode revealed that descendants of single ancestral progenitors were spatially restricted to forebrain or midbrain or hindbrain (Fig. 5a and Supplementary Fig. 6a). Such local restriction is consistent with classical single-cell labeling studies that followed cells from gastrulation to day 1 of development^37^. Notably, however, some barcodes were broadly distributed across the brain e.g. in hindbrain *and* midbrain (Fig. 5a and Supplementary Fig. 6a). This observation supports the hypothesis that some embryonic progenitors can give rise to descendants that migrate across brain regions^38^. Although barcodes were mostly regionally restricted, they were not neural cell-type restricted; single progenitors that acquired a specific barcode during late embryogenesis gave rise to descendants that mapped to multiple different clusters (Fig. 5b). In contrast to neural cells, we found more pronounced cell type restriction for non-neural cells, consistent with previous studies^23^. For example, endothelial and microglial cell lineages that shared edits with neural lineages, subsequently diverged from the neural lineages during the early barcode editing period (Supplementary Fig. 6b).

Despite the generally broad contribution of individual progenitors to multiple neural cell types, close inspection of the lineage trees also revealed divergent lineage trajectories. For example, we found that the hypothalamus/preoptic area, a brain region involved in complex behaviors such as thermoregulation, hunger and sleep, contains cell types with distinct lineage relationships. In particular, analysis of 6 barcodes across 95 cells in ZF1 indicated that there are at least two distinct neural lineages in this region: *sst3*^+^ neurons^39^ (cluster 27) were clonally related to *penkb*^+^ neurons^40^ (cluster 30), while *fezf1*^+^ neurons (cluster 20) and *hmx3a*^+^ neurons (cluster 28) were clonally related to each other (Fig. 5c, d). Inspection of the ZF1 lineage tree revealed a late barcode editing event that marked the segregation between *fezf1*^+^ neurons (cluster 20) versus *sst3*^+^ (cluster 27) and *penkb*^+^ neurons (cluster 30) (Fig. 5e). Notably, these cells were all lineage related to cluster 2 that comprised GABAergic and a small population of glutamatergic neurons in the ventral forebrain, revealing a shared common progenitor. In ZF2, 8 barcodes across 113 cells supported a similar lineage restriction (Fig. 5c and Supplementary Fig. 6c). This analysis suggests a lineage split after gastrulation between progenitors that give rise to distinct cell types in the hypothalamus/preoptic area. These results demonstrate the promise of scGESTALT to uncover the complex lineage relationships of cells with respect to cell type and position.

**Figure 6.**
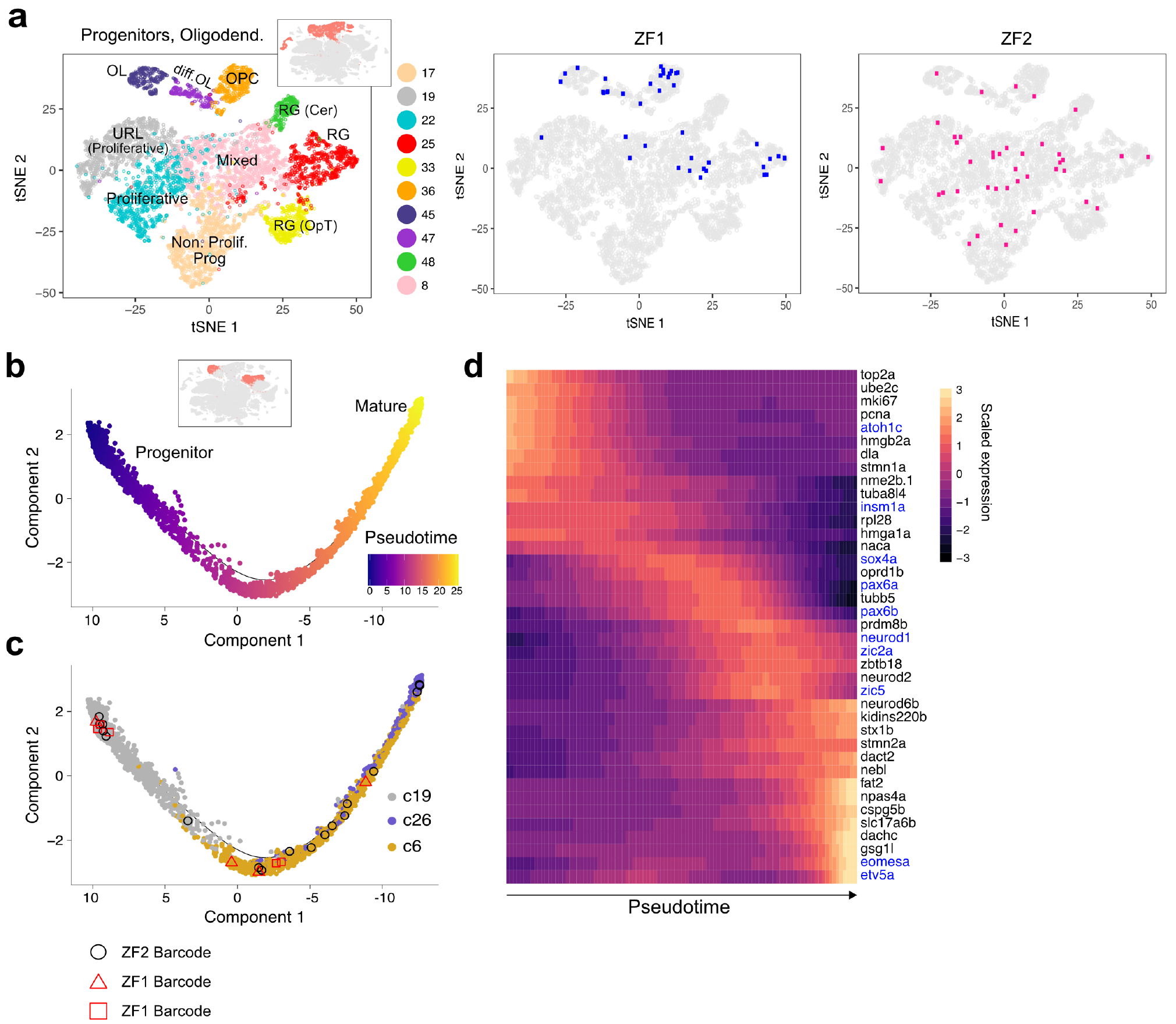
Barcodes shared between progenitor and differentiated cell types. **a**. Left, Iterative clustering of neural progenitors and oligodendrocytes. Inset highlights these clusters within the initial t-SNE plot. Right, progenitor cells from the largest barcode clone in ZF1 (blue) and ZF2 (pink) are displayed on the t-SNE plot. **b**. Trajectory of cerebellar granule cell differentiation generated with Monocle 2. Cells are colored by pseudotime. Inset highlights these clusters within the initial t-SNE plot. **c**. Cells along the trajectory are colored by cluster: 19 (progenitor); 6 and 26 (differentiated). Cells with three scGESTALT barcodes from ZF1 and ZF2 are shown to highlight barcodes found along the trajectory. **d**. Heat map of gene expression changes of selected markers during granule cell differentiation. Rows are marker genes, columns are single cells arranged in pseudotime, representative transcription factors colored in blue.

### Inheritance of edited barcodes tracks gene expression cascades during differentiation

The zebrafish brain maintains widespread neurogenic activity^41^, raising the possibility that scGESTALT could generate edited barcodes that are still shared between progenitors and differentiated cells at the time of cell isolation. Indeed, the most abundant barcodes, which comprised ~10%-26% of profiled cells, displayed broad cell type distributions (Supplementary Fig. 6d) and included 15%-28% progenitor cell types (OPCs, radial glia, intermediate progenitors) (Fig. 6a). This observation indicates that single cells marked during embryogenesis gave rise to descendants that developed both into differentiated cell types and into progenitors that maintained their capacity for neurogenesis. Although it is unknown if such late neurogenic progenitors directly gave rise to the differentiated cell types we recovered, the observed lineage relationships raised the possibility of using shared barcodes to support potential gene expression trajectories deduced from scRNA-seq data. By ordering single cells in oligodendrocyte-related clusters by gene expression signatures, we identified a trajectory from OPC to oligodendrocytes, as previously described in mouse^11,42^ (Suppl Fig. 6e-g). Similarly, cerebellar granule cell clusters followed a trajectory from *atoh1c*^+^ progenitors (cluster 19) to *pax6b*^+^ neurons (cluster 6) and then to *gsg1l*^+^ neurons (cluster 26) (Fig. 6b, c) that was accompanied by waves of gene expression changes (Fig. 6d). Strikingly, several barcodes were recovered from cells transiting through these states, raising the possibility that barcode-marked progenitor pools continue to give rise to differentiated descendants (Fig. 6c and Supplementary Fig. 6f). These results indicate the potential of scGESTALT to track gene expression trajectories during differentiation in complex multicellular systems.

## DISCUSSION

Classic studies using markers such as viral DNA barcodes or fluorescent dyes have provided fundamental insights into clonal expansion and lineage relationships during development^21,22^. The recent application of DNA editing technologies to introduce cumulative, combinatorial, permanent and heritable changes into the genome has enabled the reconstruction of lineage trees at unprecedented scales but has been limited by the lack of high-resolution cell type information and the restriction of editing to early embryogenesis^23,24,28^. Here we begin to overcome these limitations by establishing two-timepoint editing and applying scRNA-seq to identify both the identity and lineage of cells. We apply this technology, scGESTALT, to zebrafish brain development and establish its potential to simultaneously define cell types and their lineage relationships at a large scale.

scGESTALT combines the identification of cell types by scRNA-seq with the cumulative introduction of edits into a single genomic barcode (Fig. 1). The power of this approach rests on the high efficiency and diversity of barcode editing, the ubiquitous expression of the compact barcode, the ability to introduce mutations both early and late, the unequivocal recovery of the single-copy compact barcode from individual cells, the high-confidence reconstruction of lineage trees, and the simultaneous recovery of cellular transcriptomes to identify the associated cell types (Fig. 3 and 4). We foresee many immediate applications of scGESTALT in zebrafish and other model systems applying the framework introduced in this study. For example, it is now feasible to define dozens of cell types by profiling tens of thousands of cells from tissues such as spinal cord, liver, or skin using scRNA-seq and then use two-timepoint barcode editing to mark thousands of cells and reveal their lineage relationships. Variations of this approach can also be used to uncover cell type diversity and lineage relationships during tissue homeostasis and regeneration or during tumor formation and metastasis. While scGESTALT is widely applicable, several optimizations can be foreseen. First, barcode editing is still restricted to two timepoints and leads only to thousands of different barcodes. To record the full complexity of vertebrate lineage trees, future implementations will need to enable continuous editing over long time periods and generate millions or billions of differently edited barcodes. Second, the recovery of all cells and all barcodes from a single animal remains elusive. Current droplet-based approaches recover only a minority of cells, and scGESTALT currently recovers the edited barcode in fewer than 30% of transcriptomes. The comprehensive and definitive construction of lineage trees will necessitate improvements in both cell and barcode recovery. Finally, although marker genes allowed us to assign isolated cells to broadly defined regions (Fig. 2, 5), tissue dissociation results in the loss of precise spatial information. Future iterations of scGESTALT will need to identify high-resolution marker genes and create gene expression atlases to assign isolated cells to specific anatomical sites^29,43–46^.

The application of scGESTALT to brain development illustrates the potential of this approach to analyze lineage relationships in complex tissues. Our scRNA-seq analyses of the juvenile zebrafish brain identified more than 100 different cell types, provides a unique resource to identify marker genes and associated cell types, and lays the foundation to generate a complete catalogue of cell types in a vertebrate brain (Fig. 2). In combination with GESTALT, scRNA-seq generates hypotheses for potential developmental trajectories. For example, our results suggest that most descendants of an individual embryonic neural progenitor remain spatially confined but constitute multiple cell types (Fig. 4, 5). Interestingly, however, we also observed that some descendants appear to acquire a broad spatial distribution and some lineage branches separate cell types located in similar anatomical regions (Fig. 5). The inheritance of barcode edits by progenitors and differentiated cell types also raised the possibility to use such edits to help with reconstruction of developmental trajectories and the associated gene expression cascades (Fig. 6). The comprehensive understanding of the extent and emergence of such lineage restrictions and trajectories during brain development will require larger-scale lineage trees, multi-timepoint editing, and the discovery of additional cell types and marker genes.

scGESTALT lays the foundation for combining lineage recordings with single-cell measurements to reveal cellular relationships during development and disease. The finding that barcode mutations can be induced during a specific time window by an environmental signal (heat) also establishes the concept that this editing system can be rendered signal-dependent^25,26,47^. This observation opens the possibility to record endogenous or exogenous events by barcode editing: just as evolutionary history is recorded in genome sequence changes, a cell’s history might be recorded by barcode sequence edits.

## ONLINE METHODS

### Zebrafish husbandry

All vertebrate animal work was performed at the facilities of Harvard University, Faculty of Arts & Sciences (HU/FAS). This study was approved by the Harvard University/Faculty of Arts & Sciences Standing Committee on the Use of Animals in Research & Teaching under Protocol No. 25-08. The HU/FAS animal care and use program maintains full AAALAC accreditation, is assured with OLAW (A3593-01), and is currently registered with the USDA.

### Constructs for transgenesis

The GESTALT barcode transgenic vector pTol2-hspDRv7 was constructed as follows. The v7 barcode sequence^23^ was cloned into the 3’ UTR of a DsRed coding sequence under control of the heat shock (*hsp70*) promoter. This cassette was placed in a Tol2 transgenesis vector containing a cmlc2:GFP marker, which drives expression of GFP in the heart^48^.

The heat shock inducible Cas9 transgenic vector (pTol2-hsp70l:Cas9-t2A-GFP, 5xU6:sgRNA) was constructed as follows. Individual gRNAs (Supplementary Table 3) targeting sites 5-9 of the GESTALT array were cloned into five separate U6x:sgRNA (Addgene plasmids 6245-6249) plasmids, as described previously^49^. The U6x:sgRNAs were assembled into a contiguous sequence in the pGGDestTol2LC-5sgRNA vector (Addgene plasmid 6243) by Golden Gate ligation. The resulting 5xU6:sgRNA sequence was PCR amplified and ligated into the backbone of pDestTol2pA2-U6:gRNA^50^ (Addgene plasmid 63157) after the vector was first digested with Clal and Kpnl (U6:gRNA cassette of this vector was removed in the process) to generate the pDestTol2pA2-5xU6:sgRNA plasmid. The final construct was generated with multisite Gateway with p5E-hsp70l (Tol2 kit^51^), pME-Cas9-t2A-GFP (Addgene plasmid 63155), p3E-polyA (Tol2 kit) and pDestTol2pA2-5xU6:sgRNA.

Plasmids are available from Addgene - https://www.addgene.org/Alex_Schier/

### Generation of transgenic zebrafish

To generate GESTALT barcode founder fish, one-cell embryos were injected with zebrafish codon optimized Tol2 mRNA and pTol2-hspDRv7 vector. Potential founder fish were screened for GFP expression in the heart at 30 hpf and grown to adulthood. Adult founder transgenic fish were identified by outcrossing to wild type fish and screening clutches of embryos for GFP expression in the heart at 30 hpf. Single copy “heat shock GESTALT” F1 transgenics were identified using qPCR, as described previously^23,52^.

To generate inducible Cas9 founder fish, one-cell embryos were injected with Tol2 mRNA and the pTol2-hsp70l:Cas9-t2A-GFP, 5xU6:sgRNA vector. Injected embryos were heat shocked at 8 hpf and potential founder fish were screened for GFP expression at 24 hpf and grown to adulthood. F1 transgenic “inducible Cas9” fish were identified by outcrossing potential founders to wild type fish and screening clutches of embryos for whole body GFP expression after heat shock at 24 hpf.

### Two-timepoint barcode editing

sgRNAs specific to sites 1-4 of the GESTALT array were generated by in vitro transcription as previously described^23^. Single copy “heat shock GESTALT” F1 transgenic adults were crossed to “inducible Cas9” F1 transgenic adults and one-cell embryos were injected with 1.5 nl of Cas9 protein (NEB) and sgRNAs 1-4 in salt solution (8 μM Cas9, 100 ng/μl pooled sgRNAs, 50 mM KCl, 3 mM MgCl_2_, 5 mM Tris HCl pH 8.0, 0.05% phenol red). Injected embryos were first screened for GFP heart expression at 30 hpf to identify the “heat shock GESTALT” transgene These embryos were then heat shocked for 30 min at 37 C to induce Cas9 expression. Double transgenic embryos (1/4 of progeny, as expected from the genetic cross) were identified by GFP expression in the whole body.

### Preparation of GESTALT genomic DNA libraries

Genomic DNA from edited and unedited double transgenic 55 hpf embryos were extracted using the DNeasy kit (Qiagen). Samples were UMI tagged and PCR amplified using primers flanking the barcode as previously described^23^. Sequencing adapters, sample indexes and flow cell adapters were incorporated by PCR, and libraries were quantified using the NEBNext Library Quant kit (NEB). Libraries were sequenced using NextSeq 300 cycle mid output kits (Illumina).

### Whole brain inDrops

Wild type and two-timepoint edited 23-25 dpf zebrafish brains were similarly processed for inDrops single-cell transcriptome barcoding^4,53^ except that two-timepoint edited zebrafish were first heat shocked for 45 min at 37 C to induce scGESTALT barcode mRNA expression. Whole brains were dissected and dissociated using the Papain Dissociation Kit (Worthington), according to the manufacturer’s instructions with the following modifications. Brains were dissociated with 900 μl of 10 units/ml of papain in Neurobasal media (Life Technologies) and incubated at 34 C for 20-25 min with gentle agitation. Samples were then gently triturated with p1000 and p200 tips until large pieces of tissues were no longer visible. Dissociated cells were washed 2x with DPBS (Life Technologies) at 4 C and sequentially filtered through 35 μm (BD Falcon) and 20 μm (Sysmex) mesh filters. Cells were resuspended in 300-400 μl DPBS and counted using an automated Bio-Rad counter. Cells were then diluted to ~100,000 cells/ml in 18% optiprep/DPBS solution. Cells were loaded onto the inDrops device and encapsulated at a rate of 10,000-20,000 per hour. Transcriptomes were obtained for ~70% of cells introduced into the device.

### inDrops transcriptome library prep

Transcriptome libraries were prepared as previously reported^53^ with minor modifications. The product of the in vitro transcription (IVT) reaction was cleaned up using 1.3X AMPure beads (Beckman Coulter), eluted in 25 μL of RE Buffer (10 mM Tris pH7.5, 0.1 mM EDTA) and analyzed on an Agilent RNA 6000 Pico chip. 9μL of the post-IVT product was used to proceed with standard RNA-fragmentation and (untargeted) transcriptome library preparation. The remainder of the post-IVT product was left unfragmented and processed in parallel to generate scGESTALT-targeted library preps (see below).

A subset of libraries were prepared using ‘V3’ inDrops barcoded hydrogels and corresponding sequencing adapters. V3 inDrops libraries are sequenced with standard Illumina sequencing primers in which the biological read is from paired end read1, cell barcodes are from paired end read2 and index read1, and library sample index is from index read2.

### inDrops scGESTALT library prep

To generate scGESTALT libraries, inDrops samples post IVT were reverse transcribed as follows. Reactions with 5 μl IVT aRNA, 1.5 μl 50 μM random hexamer, 1 μl 10mM dNTP and 3.5 μl water were incubated at 70 C for 3 min, followed by addition of a reverse transcription mix (4 μl 5X PrimeScript buffer, 3.5 μl water, 1 μl RNase inhibitor [40U/μl], 0.5 μl PrimeScript RT enzyme). The reaction was incubated at 30 C for 10 min, 42 C for 60 min and 70 C for 15 min, and then cleaned up using 1.2X AMPure beads (Beckman Coulter) and eluted in 20 μl DS buffer (10 mM Tris pH8, 0.1 mM EDtA). scGESTALT cDNAs were PCR amplified in a two-step reaction involving: 1. GP6 and PE1S4 primers (Supplementary Table 3) and Q5 polymerase (NEB), and 2. GP12 and PE1S primers (Supplementary Table 3) and Phusion polymerase (NEB). The Q5 reaction (98C, 30s; 61C, 25s; 72C, 30s; 15 cycles) was cleaned up with 0.6X AMPure beads and eluted in 20 μl DS buffer. 8 μl of the eluate was used in the Phusion reaction (98C, 30s; 60C, 25s; 72C, 30s; 9 cycles). PCR products were once again cleaned up with 0.6X AMPure beads and eluted in 20 μl DS buffer. Finally, sequencing adapters, sample indexes, and flow cell adapters were incorporated as described for the V3 transcriptome libraries. Libraries were quantified using the NEBNext Library Quant kit (NEB).

### Sequencing inDrops libraries

inDrops V2 and V3 transcriptome libraries were sequenced using NextSeq 75 cycle high output kits. 15% PhiX spike-in was used for V2 libraries.

Sequencing parameters for V2 libraries: Read1 35 cycles, Read2 51 cycles, Index1 6 cycles. Custom sequencing primers^4^ were used.

Sequencing parameters for V3 libraries: Read1 61 cycles, Read2 14 cycles, Index1 8 cycles, Index2 8 cycles. Standard sequencing primers were used.

scGESTALT V3 libraries were sequenced using MiSeq 300 cycle kits and 20% PhiX spike-in.

Sequencing parameters: Read1 250 cycles, Read2 14 cycles, Index1 8 cycles, Index2 8 cycles. Standard sequencing primers were used.

### Bioinformatic processing of raw reads from transcriptome and scGESTALT inDrops libraries

Sequencing data (FASTQ files) were processed using the inDrops.py bioinformatics pipeline available at https://github.com/indrops/indrops. Transcriptome libraries were mapped to a zebrafish reference built from a custom GTF file and the zebrafish GRCz10 (release-86) genome assembly. Bowtie version1.1.1 was used with parameter-e 200; UMI quantification was used with parameter-u 2 (counts were ignored from UMIs split between more than 2 genes). GESTALT libraries were processed in parallel up to the mapping step with modified Trimmomatic settings (LEADING: “10“; SLIDINGWINDOW: “4:5“; MINLEN: “16“). For both scGESTALT and transcriptome libraries, error-corrected cell barcode sequences were retained for each cell to enable direct comparisons of transcript and lineage information in downstream steps. Transcriptome libraries were further processed by removing UMI counts associated with low-abundance cell barcodes. Within each biological sample UMI counts tables (transcripts x cells) were assembled.

### Cell type clustering analysis

In total, we sequenced 6,759 cells (replicate f1), 7,112 cells (replicate f2), 15,172 cells (replicate f3), 12,128 cells (replicate f4), 9,923 cells (replicate f5) and 6,026 cells (replicate f6) from whole brain samples. In addition, we sequenced 3,632 cells, 3,909 cells and 1,511 cells from manually dissected forebrain, midbrain and hindbrain regions, respectively. This resulted in a total of 66,172 single-cell transcriptomes, which were further filtered and used for clustering analysis as described below. scGESTALT libraries were prepared from whole brain replicates f3 (750 cells recovered), f5 (2,605 cells recovered) and f6 (367 cells recovered) and were designated as ZF1, ZF2 and ZF3, respectively. Clustering analysis was performed using the Seurat v1.4 R package^5,29^ as described in the tutorials (http://satijalab.org/seurat/). In brief digital gene expression matrices were column-normalized and log-transformed. Cells with fewer than 500 expressed genes or greater than 9% mitochondrial content were removed from further analysis. Variable genes (2,843 genes) were selected for principal component analysis by binning the average expression of all genes into 300 evenly sized groups, and calculating the median dispersion in each bin (parameters for MeanVarPlot function: x.low.cutoff = 0.01, x.high.cutoff = 3, y.cutoff = 0.77). The top 52 principal components were used for the first round of clustering with the Louvain modularity algorithm (FindClusters function, resolution = 2.5) to generate 63 clusters. These initial clusters were compared pairwise for differential gene expression (parameters for FindAllMarkers function: min.pct = 0.18, min.diff.pct = 0.15). Since the initial clustering contains many non-neuronal and progenitor cells, several of the top principal components were comprised of genes in those cell types. Thus, to more finely resolve transcriptional differences between neuronal clusters, select large clusters were again subjected to variable gene selection, principal components analysis, Louvain clustering and differential gene expression using the same strategy as above. This approach has been shown to uncover additional heterogeneities^42,54^. At most 12 principal components were used in these analyses. Clusters with no discernible markers or less than 10 differentially expressed genes were merged together and classified as “unassigned” clusters.

### Cell trajectory (pseudotime) analysis

Oligodendrocyte and granule cell populations were ordered in pseudotime using the Monocle 2 package^55^. The list of differentially expressed genes in each of these clusters identified by Seurat was used as input for temporal ordering in Monocle 2. The root of each trajectory was defined as the precursor (oligodendrocyte precursor cells) or progenitor (upper rhombic lip progenitors of granule cells) cell types in each of these two groups of cell populations.

### scGESTALT barcode analysis

Sequencing data from genomic DNA and inDrops scGESTALT libraries were processed with a custom pipeline (https://github.com/shendurelab/Cas9FateMapping) as previously described^23^ with the following modifications. InDrops scGESTALT reads were grouped by the inDrops cell identifiers, trimmed with the Trimmomatic software to remove low quality bases, and processed using a script designed for single-end read data. A consensus sequence was called for each single cell by jointly aligning all of its reads using the MAFFT aligner^56^. Consensus sequences were aligned to a reference sequence for the scGESTALT amplicon using the NEEDLEALL aligner^56^ with a gap open penalty of 10 and a gap extension penalty of 0.5. Aligned sequences were required to match greater than 85% of bases at non-indel positions, to have the correct PCR primer sequence at the 5' end, and to match at least 90 bases of the reference sequence. Target sites were considered edited if there was an insertion, deletion or substitution event present within 3 bases upstream of each target’s PAM site, or if a deletion spanned the site entirely. We noted that some larger inter-site deletions were misaligned or unaligned with the above parameters. These deletions were reanalyzed using the aligner from the ApE software, which searches for specified lengths of exact matching blocks of sequence, and then performs a Needleman-Wunsch alignment of the sequences between the blocks. The inDrops scGESTALT barcode for each cell was matched to its corresponding cell type (t-SNE cluster membership) assignment using the inDrops cell identifier.

To determine the stochastic nature of barcode editing, pairwise comparisons of samples were performed using cosine similarity.

### Construction of lineage trees from scGESTALT barcodes

To create the two-time-point lineage trees, scGESTALT barcodes were filtered to the editing outcomes (indels) that could only occur through the activity of Cas9 complexed to sgRNA 1 through 4 (precluding events that may start in the first 4 targets but extend into targets 5 to 9). All unique barcodes were then encoded into a paired event matrix and weights file, as described previously^23^, and were processed using PHYLIP mix with Camin-Sokal maximum parsimony^57^. In the second stage, we repeated this process for the full barcode set: each node’s descendants (barcodes that contain the identical events over the first 4 targets) were used to create a sub-tree representing the second round of editing. The original node was then replaced by this generated subtree. After the subtrees were attached, we eliminated unsupported internal branching by pruning parent-child nodes that had identical barcodes, unless this node was the junction point between the first stage node and one of its subtree members. Individual cells and their annotations were then added to the corresponding terminal barcodes. The resulting tree was converted to a JSON object, annotated with t-SNE cluster membership, and visualized with custom tools using the D3 software framework.

## ACKNOWLEDGEMENTS

For discussion and advice we thank members of the Schier lab, particularly J. Farrell. We thank the Bauer Core Facility (Harvard) and the Molecular Biology Core Facility (Dana Farber Cancer Institute) for sequencing services, and the Harvard zebrafish facility staff for technical support. This work was supported by a postdoctoral fellowship from the Canadian Institutes of Health Research to B.R., an HHMI Fellowship from the Life Sciences Research Foundation and 1K99GM121852 to D.E.W., a fellowship from the NIH/NHLBI (T32HL007312) to A.M., a Burroughs-Wellcome Fund CASI award and an Edward J Mallinckrodt Foundation grant to A.M.K., a Paul G. Allen Family Foundation grant and an NIH Director’s Pioneer Award (DP1HG007811) to J.S., a postdoctoral fellowship from the American Cancer Society to J.A.G., and NIH grants U01MH109560 and R01HD85905 to A.F.S. J.S. is an investigator of the Howard Hughes Medical Institute.

## AUTHOR CONTRIBUTIONS

B.R., J.A.G. and A.F.S. designed the study, interpreted the data and wrote the manuscript. B.R. and J.A.G. generated transgenic lines and GESTALT genomic DNA libraries. B.R. performed all experiments and data analysis, and generated scGESTALT libraries. D.E.W. generated and processed inDrops libraries. A.M. developed the scGESTALT processing pipeline and generated lineage trees. S.P. established the zebrafish neuron dissociation protocol. A.M.K. and J.S. provided resources and critical insights.

